# Open-source versatile 3D-print animal conditioning platform design for in-vivo preclinical brain imaging in awake mice and anesthetized mice and rats

**DOI:** 10.1101/2022.11.20.517296

**Authors:** Zakia Ben Youss Gironda, Tanzil Mahmud Arefin, Sawwal Qayyum, Jiangyang Zhang, Youssef Zaim Wadghiri, Leeor Alon, Omid Yaghmazadeh

**Author notes:** These authors contributed equally (Zakia Ben Youss Gironda, Tanzil Mahmud Arefin, Omid Yaghmazadeh).

## Abstract

Proper animal conditioning is a key factor in the quality and success of preclinical neuroimaging applications. We introduce an open-source easy-to-modify multi-modal 3D printable design for rodent conditioning for magnetic resonance imaging (MRI) or other imaging modalities. Our design can be used for brain imaging in anesthetized or awake mice and anesthetized rats. We show ease-of-use and reproducibility of subject conditioning with anatomical T2-weighted imaging for both mice and rats. We also demonstrate application of our design for awake fMRI in mice using both visual evoked potential and olfactory stimulation paradigms. In addition, we demonstrate that our proposed cradle design can be extended to multiple imaging modalities combining MRI, Positron emission tomography and X-ray computed tomography.

## INTRODUCTION

Preclinical animal imaging is crucial for studying and understanding biological processes *in-vivo*, disease, and function where studies are either proceeded in animal models and later on disseminated to humans or are conducted to better understand causes and effects of a disease or condition initially observed in humans [^1^]. In recent years, a growing number of preclinical Magnetic Resonance Imaging (MRI) studies have been at the research center stage, due to the multi-contrast capabilities of MRI to inform on tissue physiology and function. Functional MRI (fMRI) is currently the gold-standard technique for brain-wide imaging, enabling detection of blood-oxygenation level dependent (BOLD) related signal changes that reflect neuronal activities [^2^]. Most preclinical animal studies are limited to anesthetized animals in order to mobilize the subjects and reduce movement during imaging. However, given the non-trivial effect on brain function when using anesthetics, the translation of the information gained from these studies to animals under normal conditions remains challenging [^3,4^]. In recent years, we have seen an increase in the number of awake rodent studies allowing a more direct approach for rodent behavioral neuroscience research [^5–10^]. Importantly, awake animal studies are still in their early stages due to the challenges in proper animal setup and design of sophisticated behavioral paradigms and data analysis [^4^].

Preclinical animal imaging is challenged by suboptimal experimental solutions available to researchers, where scanners are designed for multipurpose applications aimed for broad use. This is especially problematic due to the lack of standardization among preclinical imaging vendors that makes protocols very difficult to implement in order to reliably reproduce reported studies between biomedical research groups. Notably, customized setups that are commercially available are typically proprietary to a specific vendor and can become quickly prohibitive to acquire. Alternatively, successful homemade solutions from academic research groups can be laborious to replicate without the technical expertise and resources available onsite. Second, these setups can be challenging to design in order to overcome the environment constraints associated with the MRI scanners including the limited space available within the magnet bore and the need to use non-magnetic materials while minimizing interference with the radiofrequency signal. Notably, the design of these setup is often driven by the need to facilitate accurate and reproducible positioning of the anatomical region of interest to obtain repeatable measurements. This will in turn ease high-throughput scanning for comparative studies between large animal cohorts or between imaging sessions for each individual animal during longitudinal studies. These challenges are further exacerbated by the need to ensure the proper physiological conditioning and control of the subject examined. To this effect, the design of highly customizable setups is desirable in order to accommodate awake and anesthetized animals in a repeatable and controlled manner. Moreover, the setup should provide the versatility to span across various animal types and sizes while requiring minor modifications, as well as to offer highly modular set of attachments adjacent to the animal to accommodate one or more types of stimuli. In addition to the repeatability demands, animal handling setups must be robust against imaging artifact caused throughout the scan.

Current animal conditioning platform options include commercially available products or designs developed by different research groups. Commercial designs are highly costly and lack flexibility. They are usually designed for application in a single species and for specific conditioning. The designs that we have tested have proven inefficient for both preparation time and reproducibility of the conditioning. On the other hand, designs made by research groups and reported in the literature are generally developed for specific applications and lack versatility. In addition, most of these designs are not publicly available.

In recent years the emergence of 3D printing techniques such as fused deposition modeling (FDM) and stereolithography has rapidly permeated into medicine and biomedical research [^11^]. In particular, additive manufacturing allows the generation of model shapes that cannot be created by other rapid prototyping techniques (e.g., Computerized Numerical Control (CNC) machining) with high precision, ease of manufacturing, and low costs [^12^].

Several groups have reported innovative custom 3D cradle designs customized for specific applications on anesthetized and/or awake animals (summarized in table 1.). Notably, Desai et al. initially reported an MRI compatible setup for mice with a 3D printed skull implant, exploring optogenetic stimulation performed during fMRI in awake and anesthetized animals [^5^]. Ferris et al. developed a mouse holder for fMRI in Huntington disease models with head restraining to minimize intra-scan motion [^6^]; Gilbert et al. proposed an open source cradle for various anesthetized species including mice, rats, and marmosets with support for RF electronics [^13^]; Chen et al., introduced a mouse set-up supporting somatosensory, auditory and olfactory stimulation [^7^]; Fonseca et al., developed a mouse awake animal setup for behavioral studies using 4-odor classical conditioning and visually-guided operant tasks [^8^]; Han et al., developed a mouse-holder dedicated for a cryogenic coil application on awake and behaving animals [^9^]; Sakurai et al. utilized a custom-made holder to study BOLD in mouse models of Alzheimer disease [^10^]; Donohoe et al., designed a 3D printed stereotaxic cradle for improving image quality in mouse tumor imaging [^14^]. Most of these studies are, however, not publicly available and limited to specific application (a summary of their capabilities and limitations is provided in Table 1).

**Table 1.** 3D printed designs, for small animal conditioning for imaging experiments, developed by research groups.

| Study | Printable files available online? | Easy to modify design available? | Multi species? | Applications |  |  | Head fixation mechanism |
| --- | --- | --- | --- | --- | --- | --- | --- |
|  |  |  |  | Anesthetized mice | Anesthetized rats | Awake mice |  |
| Desai *et al.*, 2011 [5] | No | No | Mice | Yes | No | Yes | Skull implant |
| Ferris *et al.*, 2014 [6] | No | No | Mice | Yes | No | Yes | Head restrainer |
| Gilbert *et al.* 2019 [13] | Yes | No | Mice/ Rats/ Marmosets | Yes | Yes | No | Head/body restrainer |
| Chen *et al.*, 2020 [7] | Yes | No | Mice | Yes | No | Yes | Skull implant |
| Fonseca *et al.*, 2020 [8] | No | No | Mice | No | No | Yes | Skull implant |
| Han *et al.*, 2019 [9] | Yes | No | Mice | No | No | Yes | Skull implant |
| Sakurai *et al.*, 2020 [10] | No | No | Mice | Yes | No | Yes | Skull implant |
| Donohoe *et al.*, 2021 [14] | No | No | Mice | Yes | No | No | Ear-bars |
| Gironda *et al.* (this study) | Yes | Yes | Mice<br>Rats | Yes | Yes | Yes | Head/body restrainer<br>Skull implant<br>Ear-bars |

Despite the proliferation of various dedicated solutions, an open source cradle that can be extended among imaging applications and modalities is still lacking. Such experimental setup should be capable of spanning across multiple modalities (Computerized Tomography (CT), Positron emission tomography (PET), MRI, etc.) while offering the flexibility to perform behavioral and functional imaging as well as to provide capabilities to attach a variety of accessories. It should also provide the ability to conduct experiments on both awake and anesthetized animals with effective restraining that is not currently widely available.

Hence, there is a crucial need for an open-source, easy-to-modify, versatile and modular, efficient, and affordable platform to perform well-controlled and repetitive experiments on small animals.

In this work, we present a solution to address these unmet needs providing an animal handling setup for multi-modal imaging in anesthetized and awake rodents. The setup is compatible with both injection-based and inhalation-based anesthesia methods and guarantees reproducible animal conditioning. We demonstrate proof-of-principle experiments conducted in the following conditions: anatomical MR imaging of anesthetized and awake mice as well as in anesthetized rats, fMRI in awake mice with successful assessment of visual evoked potential and odor stimulation, and multimodal MRI and PET/CT imaging in anesthetized and awake mice. The latest version of the computer aided design (CAD) files, including the main cradle structure, the animal head restraining using skull implant fixation, the head-fixation, the odor and light task attachments, the body elevation unit, and the anesthesia delivery system, are available online (https://github.com/omidyaghmazadeh/3D_Print_Designs/tree/main/3D_Print_Platform_for_In_Vivo_MRI_in_Mice_and_Rats).

## RESULTS

### A multi-function multi-modal cradle design for mice and rats

To answer the need for an open-source, reliable, easy to use, and easy to modify, versatile and modular 3D printable small animal conditioning set-up, we designed a system that can host both mice and rats in anesthetized state and mice in awake state, is MRI safe, is compatible with different imaging modalities and verified for fMRI in awake mice and comes with peripheral equipment for visual and odor stimulation paradigms.

The cradle was designed using Fusion 360 software (https://www.autodesk.com/products/fusion-360/). Small parts were designed for printing with a resin-based 3D printer to achieve fine resolution and were verified with one such printer (Form2, Formlabs Inc.). Larger parts were designed for filament 3D printing and were verified using PLA material (Replicator-Z18, MakerBot Ind. or Ultimaker S5, Ultimaker).

Fig. 1A illustrates the 3D design elements of the rodents conditioning system. The ‘base’ is the center-part of the system on which all other parts, depending on the application, are assembled. For example, for imaging in anesthetized mice or rats, the proper tooth/nose fixation and earbar combination should be used. Similarly, for experiments on awake mice, mouse head-fixation parts together with restraining elements are used. Add-on parts can be added to the set-up as per the need for each experiment (for example the cable/fiber holder can be used for proper positioning of the optic fiber in visual stimulation paradigms). Importantly, we suggest that a specific setup is prepared and used for each species to avoid unnecessary discomfort to the animals.

**Figure 1.**
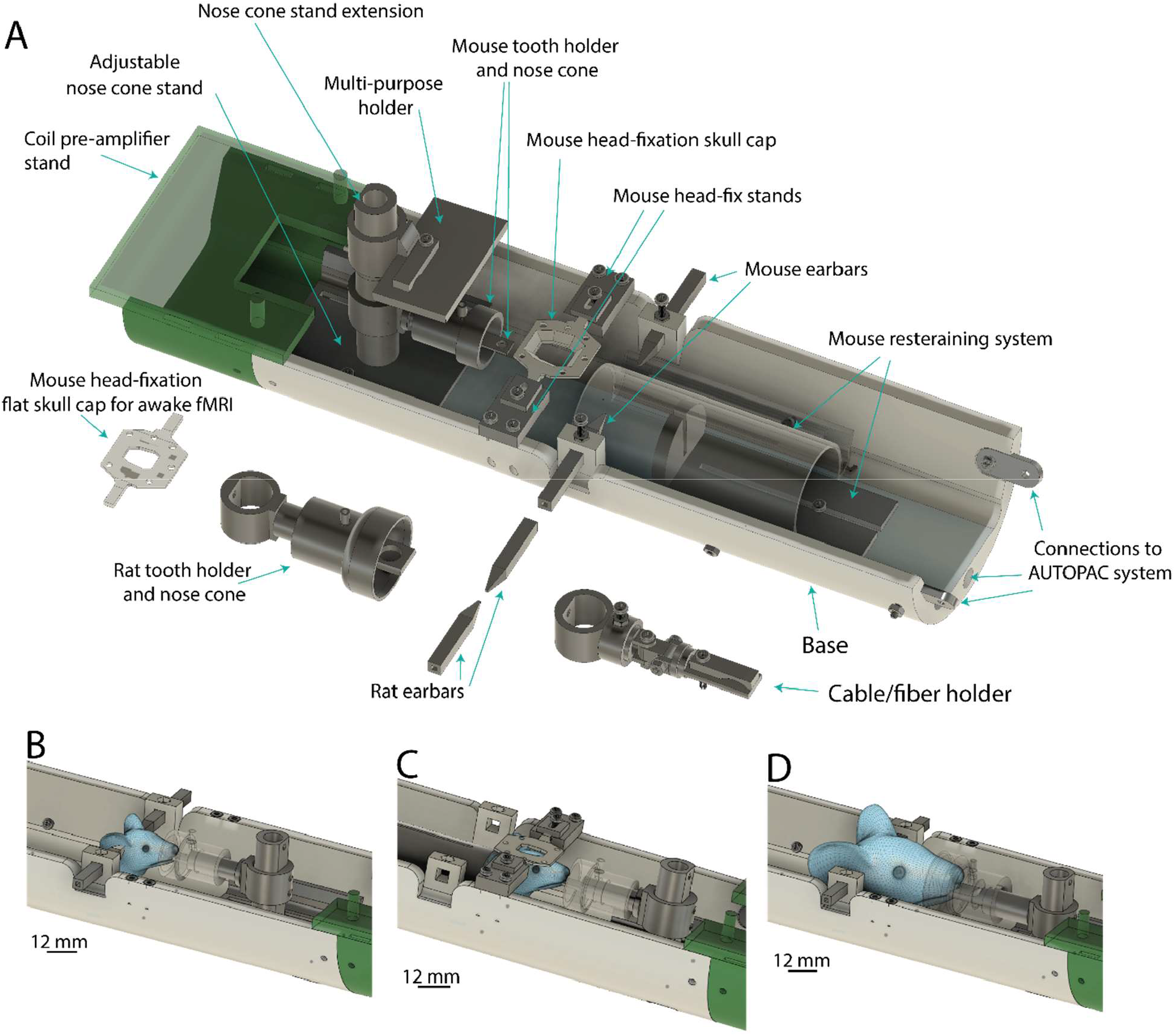
Animal conditioning platform for multimodal imaging in small animals. **A)** Schematics of the 3D-print rodent conditioning platform and its different components. **B-D)** representative graphics illustrating head fixation of the animal in different scenarios: anesthetized mice (B), awake mice (C) and anesthetized rats (D).

Our holder can accommodate anesthetized and awake mice and anesthetized rats. It is fabricated from nonmetallic MRI-compatible materials. Its base has an embedded ear-bar guiding mechanism for head-fixation of anesthetized mice and rats closely resembling the mechanism used in stereotaxic systems. The ear-bar system enables the user to achieve a reproducible head positioning while continuously delivering anesthetic agent (Fig. 1B, 1D). The base can also host a pair of removable stands for head fixation of awake mice. These stands can support matching fixtures formerly attached to the animals’ skull in a stereotaxic surgery (Fig. 1C). Importantly, our extensive experience in training and employing head-fixed mice in neuroscientific experiments (^15^) was instrumental in guiding us to successfully image awake animals. The flexible positioning of the isoflurane delivery mechanism, allows the utilization of the same setup for animals of different size (variable across sexes and ages) with minimal intervention. The modular tooth-bar stand can be extended by an add-on that permits attachment of customizable holders for experiments requiring fiber optics, cables, etc. (Fig. 1A). In awake mice experiments the nose-cone system can also be used for odor stimulation.

This animal conditioning bed is suitable for anesthetized experiment in both mice and rat using inhaled agents (e.g. isoflurane), injected agents (e.g. Urethane, Ketamine, etc.) or a combination of inhaled and injected agents (e.g. isoflurane and medetomidine). Our design supports execution of multi-modal investigations. The base of the holder can be interfaced to various imaging modalities (e.g. MRI, CT, etc) using appropriate adapter parts (Fig. 3). Remarkably, the same head-fixation system for awake mice can be used in several neuroscientific experiments such as calcium imaging. These characteristics facilitate the design and execution of multi-modal investigations.

Tapping 3D-printed plastic materials during assembly often relies on the quality of the printed part and the experience of the assembler, and the tapped parts tend to wear off over time and compromise the robustness of the whole structure especially during extensive usage. To this effect, we opted for a tap-free approach in order to secure the 3D printed parts using nylon nuts glued to the printed parts where needed (Supp. Fig. 1).

We examined the holder in various conditions and for different applications to validate its proper functionality as follows:

### Morphological brain imaging of mice and rats

Brain morphological images were acquired using T2 RARE sequences (see Methods) from anesthetized mice and rats (n=5 and n=3), and awake mice (n=5), respectively (Fig. 2A-C). The animal preparation time (excluding the anesthetization in the isoflurane chamber) was less than 2 minutes per animal. Fig. 2A-C(i) illustrate the animal cradle with mice and rat in anesthetized and awake states prepared for imaging. Fig. 2A-C(ii), shows reproducibility of slice images in animal brain with minimal preparation time (~ 2minutes per animal), with no apparent imaging artifacts due to the set-up. This reflects the consistency in alignment and positioning of the animals’ head across subjects which is a prerequisite for reliable and reproducible MR data acquisition, processing, and interpretation. Additionally, the resulting anatomical images were successfully adopted to pre-registered brain atlas images (Fig. 2D), providing another demonstration of the stability of the animal conditioning using the cradle. Respiration motion artifact during MRI acquisition can cause ghosting effects, however, our imaging results did not show ghosting or other forms of artifacts. In addition, applying motion assessment in the images acquired for rats (on the 10 repetitions of each slice), we observed minimal respiration-induced motion artifacts (Supp. Fig. 2). This demonstrates the resilience of the set-up to motion while minimizing the preparation time required for each animal, thus providing a robust solution for animal conditioning with reproducible results.

**Figure 2.**
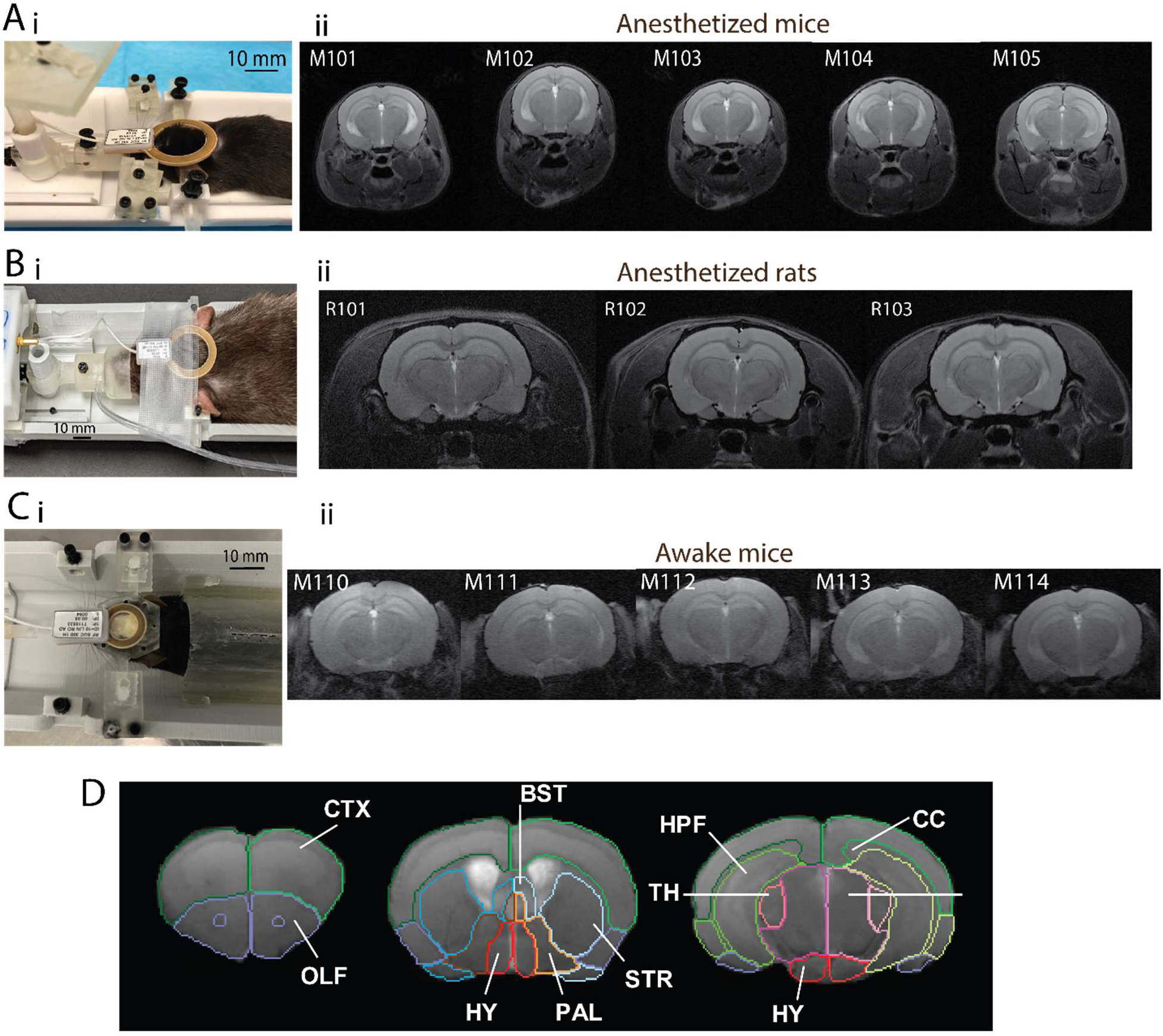
Animal conditioning and magnetic resonance imaging in anesthetized and awake mice and anesthetized rats. **A)** i: The rodent cradle with anesthetized mouse preparation using the tooth-bar/ear-bar combination for head fixation, and nose-cone for isoflurane delivery. The picture shows the final animal preparation with a receive-only coil (ID=20 mm) placed on top of the animal’s head for image acquisition. ii: T2 Rare images of anesthetized mice (n=5) prepared as shown in A (TR=2500-ms, TE=33-ms, Turbo-factor=8, FOV=20 mm x 20 mm, Matrix size=256 × 256 resulting in 78 µm × 78 µm in-plane resolution, 19 slices of 800 µm thickness, with a total scan time of 13 min). **B)** i: The rodent cradle with anesthetized rat preparation using the rat-size tooth-bar/ear-bar combination for head fixation, and nose-cone for isoflurane delivery. The picture shows the final animal preparation with a Rx coil (ID=20 mm) placed on top of the animal’s head for image acquisition. ii: T2 Rare images of anesthetized rats (n=3) prepared as shown in A (TR=2500-ms, TE=33-ms, Turbo-factor=8, FOV=35 mm x 35 mm, Matrix size=256 × 256 resulting in 137 µm x 137 µm in-plane resolution, 19 slices of 800 µm thickness, with a total scan time of 13 min). **C)** i: Example of an awake mice conditioned on the platform. The preparation consisted of attaching the head-fixed implant (installed during a pre-experiment surgery) to the corresponding joint pieces. The animals were constrained using a restraining component designed to a snug fit to minimize body movement. A receive coil (ID=10 mm) is placed on top of the animal’s implant for image acquisition. ii: T2 Rare images of awake mice (n=5) with the following parameters: TR=2516 ms, TE=35 ms, Rare-factor=8, FOV=30 mm x 9 mm, Matrix size=256 × 256 resulting into 117 µm x 35 µm in-plane resolution, 18 slices of 800 µm thickness, with total scan time 2min and 41 sec. D) Identification of brain structural labels from T2 images for the example of an anesthetized mouse as an indirect way to demonstrate the stability of the animal conditioning during the imaging experiment. Mouse brain atlas (^16^) were coregistered into representative T2 image of one of the awake mice brain. Brain region annotations are **CTX**: Cortex, **OLF**: Olfactory area, **HPF**: Hippocampus, **STR**: Striatum, **TH**: Thalamus, **PAL**: Pallidum, **HY**: Hypothalamus, **MB**: Midbrain, and **CC**: Corpus callosum.

### Animal preparation for awake experiments

For experiments in the awake state, animals undergo a surgery for implantation of a head-fixation cap on their skulls (see Methods). After post-surgery recovery time (≥ 5 days) animals are trained for head-fixation. The first 5 days of training occur outside the MRI bore with an incremental timing to slowly habituate the animal. Animals are then trained for 5 additional days inside the scanner with incremental total to gradually habituate to the environment. Dummy scan (identical to actual fMRI scans with parameters described further below) times are also incrementally incorporated into the training (See Methods).

### Multimodal imaging in anesthetized and awake mice

To demonstrate the adaptability of our cradle design to different imaging modalities we designed an adapter component (Fig. 3Ai and 3Bi) to connect the cradle to the holding system of a PET/CT scanner (Inveon, Siemens). The same setup used for MRI, could be transferred to the PET/CT and connected via the adapter for PET/CT image acquisition. The orientation of the mouse is kept the same as in MRI experiments to ease the off-line software co-registration between MRI and PET/CT data. The adapter is designed (with a sled configuration) to fit with the Siemens Inveon PET/CT scanner. Figure 3Ai shows the preparation of an anesthetized mouse for PET/CT acquisition including the cradle connected to the adapter. Similarly, Fig. 3Bi illustrates an awake mouse prepared for PET/CT acquisition (note that there is no isoflurane delivery for the awake animal). Both animals were imaged first using MRI, then transferred shortly after, and prepped for PET/CT acquisition. The preparation for the PET/CT acquisition consisted of connecting the PET/CT adapter and transferring the holder the PET/CT system where the adapter was connected to its motion bed. The set-up is then moved inside the scanner for PET/CT acquisition. Each animal was injected with Fludeoxyglucose (FDG) dose (200 mCi) and we dynamically acquired the PET data. In the case of the anesthetized mouse, the body temperature of the animals was maintained by using an air heater supply during the scan. Both anesthetized mouse and awake mouse were imaged under similar conditions and the same PET/CT acquisition parameters were applied. Fig. 3Aii, 3Bii, show the resulting PET and CT images from the awake and anesthetized animals using the following PET acquisition parameters: Emission, 1800 sec, axial length = 127 mm; and CT acquisition parameters: attenuation scan, 528 sec, effective pixel size= 99.89 um, transaxial FOV=51.15 mm, Axial FOV=79.92 mm, exposure settings (80 KV, 500 µA, filter 0.5 mm, exposure time 180 ms).

**Figure 3.**
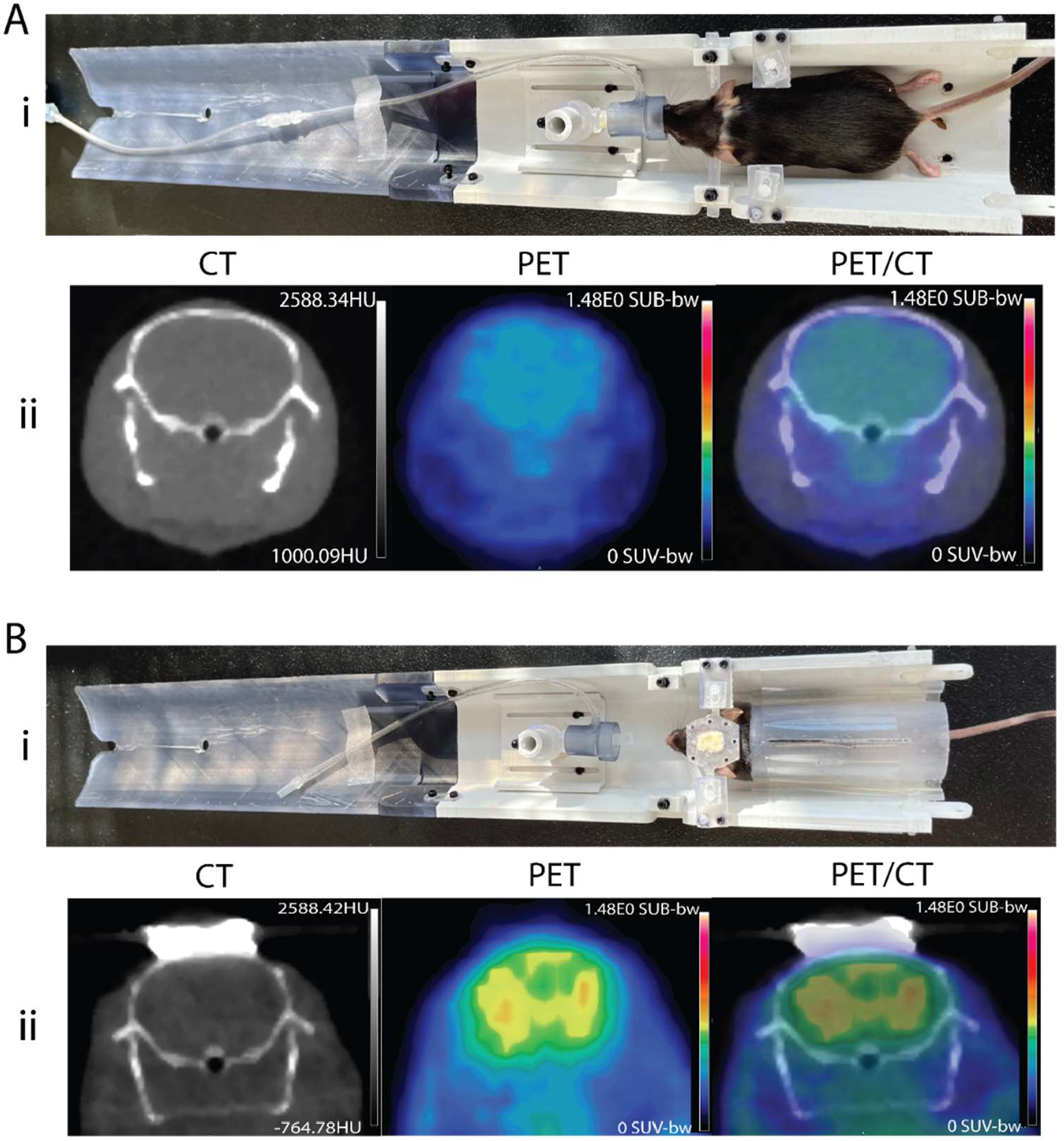
Multi-modal imaging in anesthetized and awake mice. **A)** i: The picture shows the PET/CT adapter connected to the animal conditioning platform with an anesthetized mouse. ii: CT, PET, and PET/CT overlay images obtained from the anesthetized mouse using the following PET acquisition parameters: Emission, 1800 sec, axial length=127 mm and CT acquisition parameters: attenuation scan, 528 sec, effective pixel size= 99.89 um, transaxial FOV=51.15 mm, axial FOV=79.92 mm, exposure settings (80 Kv, 500 uA, filter 0.5 mm, exposure 180 ms). **B)** i: The picture shows a similar preparation set-up an in A but for an awake mouse. ii: CT, PET, and PET/CT overlay images obtained from the awake mouse using the following PET acquisition parameters: Emission, 1800 sec, axial length=127 mm and CT acquisition parameters: attenuation scan, 528 sec, effective pixel size= 99.89 um, transaxial FOV=51.15 mm, axial FOV=79.92 mm, exposure settings (80 Kv, 500 uA, filter 0.5 mm, exposure 180 ms).

### fMRI experiments in head-fixed awake mice

We demonstrated the application of our animal conditioning system for fMRI experiments in awake mice. We performed resting-state fMRI (rs-fMRI) as well as fMRI responses to visual evoked potential (VEP) and odor stimulation paradigms.

Fig. 4A shows the experimental set-up for the fMRI experiments in VEP (i) and olfactory stimulation (ii) together with the applied imaging/stimulation paradigms. rs-fMRI is performed in both of these experiment prior to application of sensory stimulation. Fig. 4B illustrates the image quality assessment of acquired data using rs-fMRI, VEP fMRI, and odor stimulation fMRI (including temporal signal-to-noise ratio (tSNR), frame displacement (FD), and brain activity Z-score) confirming the capability of the set-up for fMRI experiments in awake mice.

**Figure 4.**
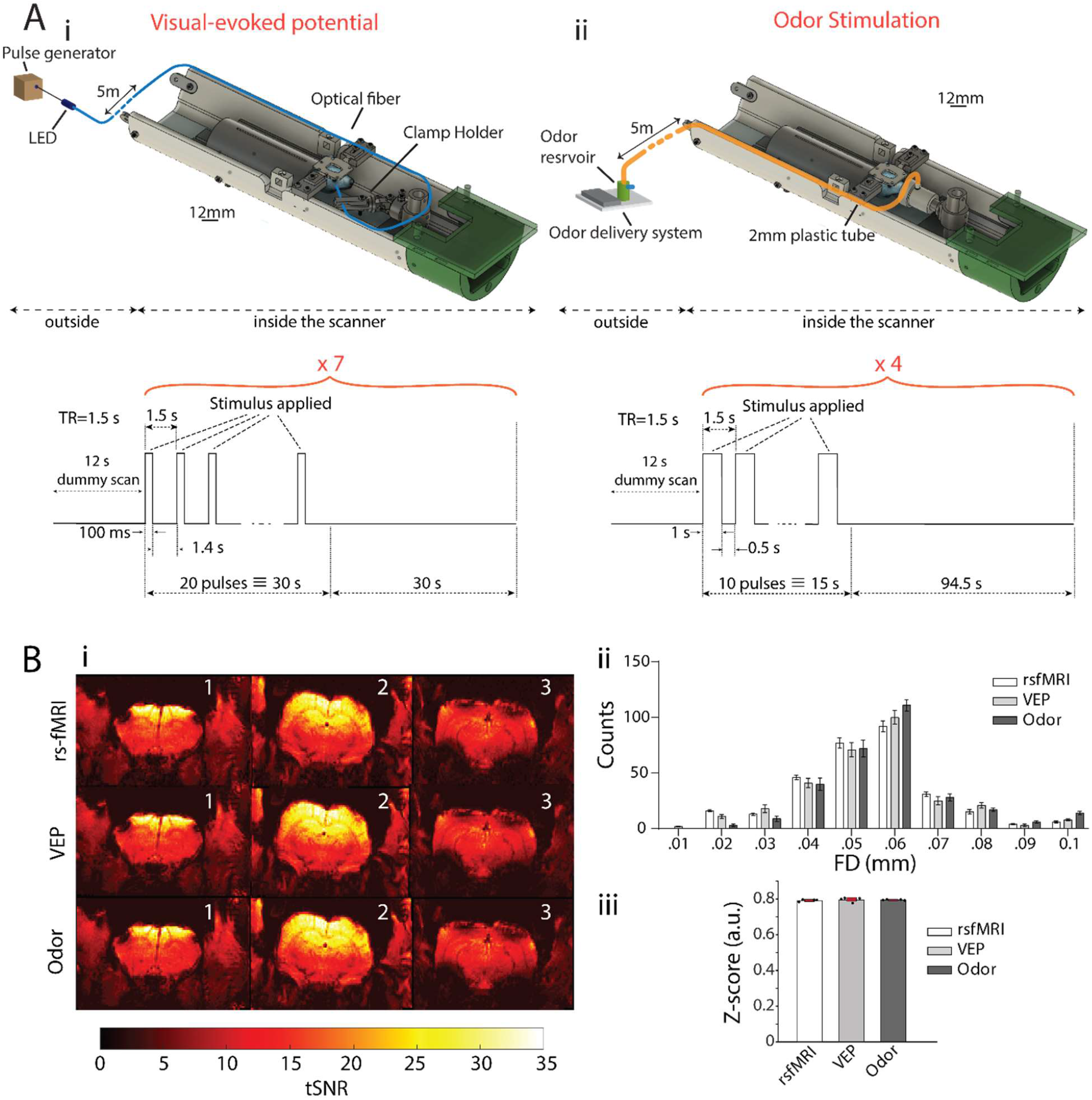
Functional magnetic resonance imaging in awake mice: set-up and quality controls. **A)** The experimental set-up and the timing paradigm used in VEP (i) and olfactory stimulation (ii) fMRI experiment. **B)** Image quality assessment of the awake mouse brain fMRI acquired using resting-state functional MRI (rsfMRI), visual evoked potential (VEP), and odor stimulation; i: Temporal signal-to-noise ratio (tSNR). ii: Frame displacement (FD), shows no apparent frame-wise displacement across scans and iii: Brain activity – Z-score, demonstrates no aberrant activation of the brain due to stimulation in awake mice.

### Resting-state mouse brain network is preserved in awake state

Using bi-lateral retrosplenial cortex (RSP) as the seed region, we tested whether the cradle could maintain the resting-state during awake condition and identify the default mode network (DMN) network in mouse brain. Fig. 5Ai shows the mouse brain cortical atlas and seed region (RSP) overlaid on template T2 image. We perceived that RSP – the core hub of the DMN network is positively correlated with the medial and caudal anterior cingulate area (ACA), posterior parietal association area (PTLp), temporal association area (TEa), sensori-motor area (SS/MO), and visual area (VIS) of the cortex, as well as hippocampus (HPF), thalamus (TH) and midbrain (MB) (Fig. 5Aii), similar to earlier reports in rodents (^17–20^) and human (^21^). Preservation of the DMN in awake mouse brain rs-fMRI suggests that our customized 3D printed cradle is highly capable of maintaining stable position and animal physiological conditions during the scan.

**Figure 5.**
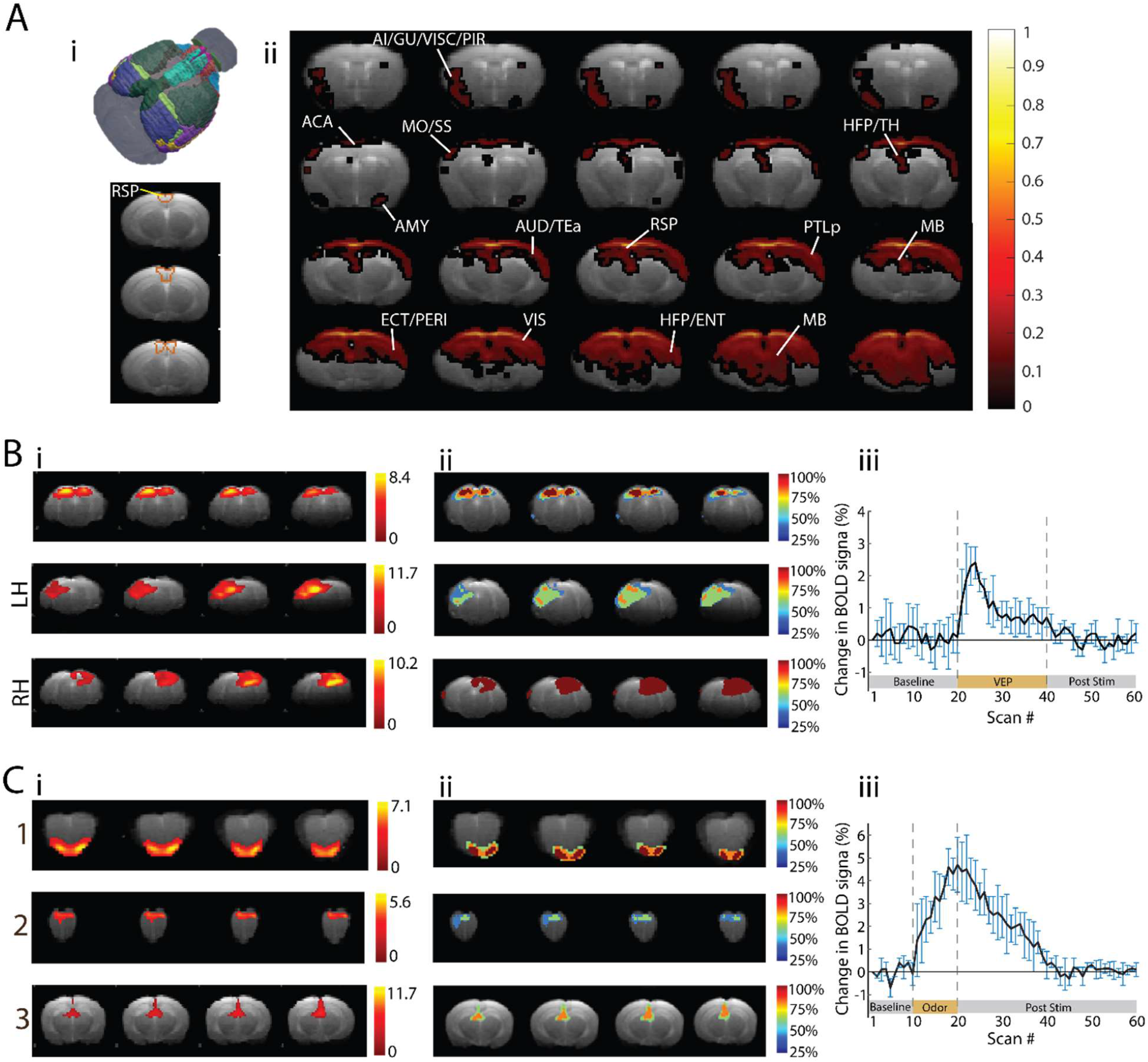
fMRI responses in awake mice. **A)** Patterns of default mode network (DMN) in awake mouse brain revealed by resting state functional MRI; i: 3D mouse brain isocortex atlas and 2D representation of the three consecutive axial slices of the bi-lateral retrosplenial cortex (RSP) – core hub of the DMN. ii: Group average functional connectivity map from RSP region demonstrated strong coherent fluctuations of the BOLD signal with the cortical regions including rostromedial ACA, MO, SS, TEa, AI, GU, VISC, PIR, ECT, PERI and VIS areas, as well as subcortical regions, such as, HPF, AMY, TH, and MB. **B)** i: Representative axial slices of the activation patterns of bi-lateral visual cortex (VIS) including superior colliculus (SC) during Visual evoked potential (VEP) fMRI. ii: Incidence maps of the identified functional clusters showing the percentage of activated voxels among subjects. iii: Percentages of changes in BOLD signal (mean ± s.d.) in awake mouse visual cortex induced by VEP-fMRI (n=4 mice). **C)** Odor stimulation induced activation of the bi-lateral; i: Olfactory area (OLF; 1), Orbital cortex (OFC; 2), and Mediodorsal nucleus of the thalamus (3) and ii: the corresponding incidence maps of the functional clusters are shown in the right panel. Iii: Percentages of changes in BOLD signal (mean ± s.d.) in awake mouse olfactory area induced by odor stimulation (n=4 mice).

### Identification of the functional clusters activated during non-tactile stimulation

To identify the elementary functional clusters due to visual and olfactory stimulation in awake mice, we applied data driven spatial group independent analysis (ICA, 20 components). We identified only one artifactual component from olfactory stimulation paradigm related to cerebrospinal fluid or vascular activation. As expected, VEP fMRI induced significant activation in the mouse brain bi-lateral visual area (VIS) including superior colliculus (SC) and the dorsal part of the lateral geniculate complex (LGd) (Fig. 5Bi) and olfactory stimulation regime significantly activated the olfactory area (OLF) as well as orbito-frontal cortex (OFC) and medio-dorsal nuclei of thalamus (MD) (Fig. 5Bii). We further justified the reliability and reproducibility of the functional clusters in the primary activation sites (VIS and OLF) during visual and olfactory stimulation paradigms by computing spatial, back-reconstructed, individual subject components using a spatial-temporal regression approach and created incidence maps (^18^). We observed that 75%-100% voxels in the primary activation area were consistently activated among the subjects, indicating high reliability and reproducibility of the stimulation-induced functional activity in the brain using the cradle. In addition, we examined the percentage of changes in the BOLD signal from baseline to stimulation timepoints (VEP and olfactory). We observed significant changes in the BOLD signal in the VIS – SC area immediately after the onset of VEP, reaching its peak (2.4%) around 4.5s after the first pulse with an average BOLD amplitude of 0.85±0.3% over the following stimulation timepoints and eventually returns to baseline during the post-stimulation period (mean: 0.1%) (Fig. 5Biii). During the olfactory stimulation, significant BOLD activation was observed in the OLF area (peak: 4.7%) with an average activation of 3.16±1.1% over the following stimulation timepoints and further returning to baseline during the post-stimulation period (Fig. 5Ciii).

## DISCUSSION

We developed a 3D-print design to accommodate mice and rats for imaging experiments. We demonstrated its ease-of-use, fast, repeatable and accurate preparation using anatomical imaging of multiple anesthetized mice and rats and awake mice. In addition, we showed that this platform, together with specific accompanying add-ons, can be successfully used for complex imaging experiments such as awake fMRI. Furthermore, we demonstrated its capability to adapt to different imaging modalities by a dual MRI-PET/CT imaging experiment.

Ready to print ‘.stl’ files for all parts are provided online. In addition we provide the original design (‘.f3d’ file format developed using the Autodesk Fusion360 software) to make it easier for users to modify the design as per their needs, whether they would like to add new parts or adjust the existing ones.

When designing this system, we took a tap-free strategy. Tapping for screws is a time consuming process which might end up in low quality results in some printed plastic materials and they usually wear off over time. Instead, we accommodated appropriate room for hosting commercially available high-quality nylon nuts wherever a screw is applied.

### Laboratory-based 3D printing

Utilizing 3D printing capabilities in a laboratory setting has proven to be a reliable and robust modality with respects to designing specific use cases in a multitude of projects. Primarily, having a 3D printer in house, can allow a lab to address specific design requirements not met by commercial suppliers or in a much affordable manner. With growing commercial printing services across the world, research laboratories can still benefit from 3D printing with reasonably affordable costs. Utilization of 3D printers has rapidly gained traction in the clinical space [^22^], however, its utility in the preclinical space is just as appreciated spanning a range of applications from 3D bio-printing to more device-oriented usages.

Within the scope of 3D printing, there are generally two major forms of desktop form printers readily accessible to the average consumer, fused deposition modeling (FDM) and stereolithography (SLA), both coming with their respective strength and weaknesses, which are beyond the context of this narrative. In summary, FDM printing (which uses thermoplastic filament as source material) have a simple design and provide a vast scalability while SLA printing (which is resin-based) provides a significantly better print resolution and overall better structural print quality. For printing the prototypes and final set-ups used in this study we employed a FDM printer (Replicator-Z18, MakerBot Ind. or Ultimaker S5, Ultimaker) for printing the larger parts (e.g., the base, the adjustable stand, etc.) and a SLA unit (Form2, Formlabs) for printing parts that require better resolution (e.g., the head-fixing cap, the earbars, etc.). One who wishes to reprint this design should have in mind that printing with different printers might need few rounds of tweaking to achieve satisfactory outcome.

### Visuo-thalamic and olfactory-thalamic responses during non-tactile stimulation

Thalamus is a salient hub for multiple sensory functions including visual, auditory, and olfaction. Visual information in the brain is processed through a series of interactions between the retina, thalamus and the visual cortex that are preserved across species (^23^). Using MRI, the projection of optic nerves to the geniculate complex of the dorsal and ventral thalamus (LGd and GENv) through the optic chiasm that enroute to the superior colliculus (SC) and visual cortex (VIS) has previously been reported (^24,25^). We perceived significant BOLD activations in mouse visuo-thalamic network highlighting functional interactions of the visual cortex with the geniculate complex of the thalamus and SC.

In olfaction, mediodorsal nucleus of the thalamus (MD) receives input from the piriform cortex (PIR) that in turn, projects to the orbito-frontal cortex (OFC) forming a transthalamic pathway in the brain (^26^). During odor stimulation in awake state of mice, olfactory region showed significant interactions with the MD and OFC. These findings indicate that our custom designed cradle can reproduce the complex functional interactions among the brain regions during the non-tactile stimulation fMRI.

### Subject population of fMRI studies

Usually when fMRI is applied to monitor the neural signature of a paradigm in rodents, 10-20 animals are used to ensure reproducibility of the data. In our study, however, as we only aimed to demonstrate the proof of principle for application of our animal conditioning set-up in fMRI assessment in awake mice using well-established stimulation paradigms (i.e., VEP and odor stimulation) we limited the cohorts to four animals. Our results, showing consistent outcomes, validates this decision and confirmed that it was not necessary to engage a larger number of animals.

### Potential usage for experiments with awake rats

We have tested our animal conditioning platform with anesthetized and awake mice and anesthetized rats. It is however possible to adapt the design, with small modifications, so that it could be used for experimentation with restrained awake rats. The head-fixation should be modified to support higher mechanical strength against animals’ head movements and the restraining mechanism should also be adapted to the bigger size of rats’ body.

### Add-ons

The presented design is versatile in the sense that it can be adapted to the needs of the desired experiment. Add-ons are the key elements to support such versatility. Although some add-on parts are presented in this study, there are many more possibilities not covered. For example, a temperature feedback-controlled electrical heating pad can be added to the cradle for better control of the animal’s body temperature in anesthetized animal experiments.

## Acknowledgements

Authors thank Mihály Vöröslakos whose designs have been an inspiring source for some of the elements in the presented study. Authors thank members of the BuzsakiLab (buzsakilab.com), and labs of Drs. Jiangyang Zhang, Youssef Z. Wadghiri and Leeor Alon and the team of the preclinical imaging core at the NYU Grossman School of Medicine for their help and feedback on different aspects during the evolution of this work. This work was supported by NIH grant #1R01NS113782-01A1 and TL1 postdoctoral fellowship # 2TL1TR001447-06A1 to OY. All the imaging experiments were performed at the NYU Langone Health Preclinical Imaging Laboratory, supported by the NIH/SIG #1S10OD018337-01, the Laura and Isaac Perlmutter Cancer Center Support Grant #NIH/NCI 5P30CA016087, and the NIBIB Biomedical Technology Resource Center Grant NIH #P41 EB017183 as well as the NYU CTSA grant #UL1 TR000038 from the National Center for Advancing Translational Sciences, NIH.

## Author contributions

O.Y. and L.A. conceived the project and O.Y. coordinated its execution. O.Y. designed the platform with input from Z.B.G., L.A., S.Q. and Y.Z.W. O.Y., Z.B.G., T.M.A., L.A., Y.Z.W., and J.Z. designed the experiments. O.Y., Z.B.G. and T.M.A. performed the experiments and analyzed the data. O.Y., Z.B.G, T.M.A and L.A. wrote the paper and all authors participated in its revision. Authors declare no competing interest.

## METHODS

### Animals

#### Mice

adult wild-type C57BL/6JxFVB mice (25-33 gr) were obtained from Charles River Laboratory. Mice were kept in cages in a 12hr regular cycle vivarium room dedicated to mice in up to five-occupancy cages.

#### Rats

adult Long-Evans rats (400-500 gr) were obtained from Charles River Laboratory and kept in a 12hr regular cycle vivarium room dedicated to rats in double-occupancy cages.

In all experiments, each animal served as its own control, no randomization or blinding was employed. No prior experimentation had been performed on the animals.

Animals with chronic head-caps were moved to single occupancy cages to avoid both social conflicts between animals and damage to the implanted materials. All experiments were conducted in accordance with the Institutional Animal Care and Use Committee (IACUC) of New York University Medical Center.

### MR imaging in anesthetized mice and rats

#### Stereotaxic fixation

Both mice and rats were first anesthetized in an induction box supplied with a mixture of 0.5-2% isoflurane and air. Animal’s head was stereotaxically fixed using the appropriate tooth bar and ear bars to avoid possible head motion during the scan. The nose cone was attached to the isoflurane vaporizer system (EZ-155, World Precision Instruments Inc.) by a single 2 mm diameter plastic tube and a male Luer connector. Animal’s body temperature and respiration rate were monitored throughout the scan using a rectal temperature probe (SA Instruments) and a pressure sensitive respiration pad (SA Instruments), respectively. Animal body temperature was kept constant (36° to 38° C) using a heating pad consisting of tubes filled with hot water connected to a water heater circulating system (Thermo Fisher). An 86 mm inner diameter birdcage coil (Bruker Corp.) - was used in transmit mode, in combination with a loop coil (ID=20 mm; Bruker Corp.) for signal acquisition in RX mode. The Rx loop coil was placed on top of the animal head as close as possible but without touching it and secured in place using surgical tapes. The cradle was then inserted inside the scanner using Bruker Autopac system for brain imaging.

#### MR data acquisition

For all animals, firstly, a pilot scan was acquired using Bruker localizer protocol (3D FLASH sequence) to ensure proper positioning of the animal.

High-resolution morphological images covering the whole animal brain were acquired using T_2_-weighted Turbo rapid acquisition with relaxation enhancement (RARE) sequence using the parameters listed in Table 2.

**Table 2.** Parameters used in imaging of anesthetized mice and rats.

| Parameters | Mouse | Rat |
| --- | --- | --- |
| TE/TR (ms) | 33/2500 | 33/2500 |
| Averages | 10 | 10 |
| RARE factor | 8 | 8 |
| Field of view (FOV) (mm) | 20 x 20 | 30 x 30 |
| Matrix size | 256 x 256 | 256 x 256 |
| No. of axial slices | 19 | 19 |
| Slice thickness (mm) | 0.8 | 0.8 |
| Scan time | 13 mins | 22 mins |

### Head-fixation cap implantation surgery for MR imaging of awake mice

A mouse head-fixation set-up with a matching head-mount cap was incorporated in the cradle design for experimentation with awake mice. Mice underwent a short surgery for attachment of the head-cap to their skulls.

During the surgery, mice were kept anesthetized under a steady stream of isoflurane (2%). The rectal temperature was kept constant at 36–37 °C with a DC temperature controller (TCAT-2, Physitemp LLC, Clifton, NJ) and stages of anesthesia were maintained by confirming the lack of vibrissae movements and nociceptive reflex. Skin of the head was shaved, and cleaned by three successive application of povione-iodine surgical scrub solution and alcohol pads. The skin was retracted after a medio-sagittal incision, and the bone surface was cleaned with hydrogen peroxide (2%) and let to dry. Metabond (Parkell Inc, Edgewood, NY) was applied on the skull surface and the custom-designed, 3D-printed plastic head-cap was then attached to animal’s skull using dental cement. Animals were allowed to recover from surgery (for at least 4 days) before conducting any experiments.

### Mice habituation for experiment with Head-fixation

Before starting the experiments, animals were trained to habituate with head-fixation for incremental duration from 5min to up to 2hrs. The training was first conducted outside of the scanner over 5 days (Day1: 1min, 5min, 15min head-fixation with 20min rest times in between; Day2: 5min, 15min, 30min head-fixation with 30min rest times in between; Day3: 5min, 30min, 60min head-fixation with 60min rest times in between; Day4: 10min, 30min, 90min head-fixation with 60min rest times in between; Day5: 10min, 30min, 120min head-fixation with 60min rest times in between). They were then trained inside the scanner over additional 5 days including incremental dummy scan times (Day1: 1min, 5min, 15min head-fixation with 20min rest times in between; Day2: 5min, 5min (with dummy scan), 30min (with dummy scan) head-fixation with 30min rest times in between; Day3: 10min, 30min (with dummy scan), 60min (with dummy scan) head-fixation with 60min rest times in between; Day4: 10min, 30min (with dummy scan), 90min (with dummy scan) head-fixation with 60min rest times in between; Day5: 10min, 30min (with dummy scan), 120min (with dummy scan) head-fixation with 60min rest times in between). Dummy scans parameters were identical to those applied in the fMRI experiments as described in Table 4.

### MR imaging in head-fixed awake mice

Trained mice for head-fixation were conditioned inside the cradle and their head-fixation cap was secured using the specific holders. The conditioning process was relatively very fast, and due to the matching between the head-cap and the corresponding holders, the resulting head positioning was highly stable across animals. After securing the mouse head to the platform, the animal’s body was covered with a 3D printed constraining element to minimize motion and stress during the imaging. The laser centering was used along with the scanner’s bed system to insert the cradle carrying the animal into the MRI bore. A pilot scan was first acquired using Bruker localizer protocol (3D FLASH sequence) to ensure proper positioning of the animal.

#### T2 imaging

Similar to the anesthetized animals, imaging of awake head-fixed animals started with performing a FLASH sequence to verify animal’s head centering, followed by morphological T2-weighted RARE sequence (with two averages with turbo-factor of 8) using the parameters listed in Table 3.

**Table 3.** Parameters used in imaging of awake mice.

| Parameters | Values |
| --- | --- |
| TE/TR (ms) | 35/2516 |
| Field of view (FOV) (mm) | 30 x 9 |
| Matrix size | 256 x 256 |
| No. of axial slices | 19 |
| Slice thickness (mm) | 0.8 |
| Planar resolution (mm) | 0.117 x 0.035 |
| Scan time | 2mins 41sec |

### fMRI experiments in head-fixed awake mice

To test the feasibility and efficacy of the cradle in awake animal MR imaging, we conducted multiple fMRI experiments including resting-state fMRI (rs-fMRI), non-tactile (visual and olfactory) sensory stimulations. Imaging sessions were initiated by a pilot scan using Bruker localizer protocol (3D FLASH sequence) to ensure proper positioning of the animal.

*fMRI data acquisition*: Both the rs-fMRI and stimulation induced fMRI (visual evoked potential (VEP) and olfactory stimulation) data were acquired using T2*-weighted single shot gradient echo – echo planar imaging (GE-EPI) sequence (^18,27^) with the imaging parameters listed in Table 4.

**Table 4.** Imaging parameters used for fMRI experiment in awake mice.

| Parameters | Values |
| --- | --- |
| TE/TR (ms) | 13.7/1500 |
| Field of view (FOV) (mm) | 30 x 9 |
| Matrix size | 256 x 256 |
| No. of axial slices | 14 |
| Slice thickness (mm) | 0.8 |
| Planar resolution (mm) | 0.234 x 0.14 |
| Scan time | 7mins 30sec |

Rectal temperature and respiratory rate were monitored throughout the experiment using a MRI compatible rectal probe, and a pressure sensitive pad placed underneath the mouse abdomen.

#### Visual evoked potential stimulation

VEP was introduced using a fiber-coupled light source placed outside of the scanner room. Seven consecutive sets of 20 short (100 ms ON, 1400 ms OFF) white light pulses (total of 30 s) followed by 30s OFF period (Fig. 4Ai) was introduced to animal’s right eye through the tip of the fixed fiber (held by the holder component of the 3D design).

#### Olfactory stimulation

Mouse brain olfactory network was stimulated using a custom-made odor delivery system (Suppl. Fig. 3) consisting of commercially available low-cost parts including an air pump and two normally-off valves. A constant shallow air flow is delivered to animal’s nose through the mice nose-cone component (which is also used for delivery of the anesthetic agent in experiments with isoflurane-anesthetized animals). Four consecutive sets of ten (1s ON, 0.5s OFF) electrical pulses (total of 15s) followed by 94.5s OFF period (Fig. 4Aii) were applied to open the valves and switch the piping circuit to conduct the air flow through a container with an odor agent (17% isopropanol alcohol in our experiments). As the same air pump is used for both cases, the total air flow stays the same for presence and absence of the odor agent.

#### Data quality check

Quality of the awake fMRI datasets was vigorously checked by computing the temporal signal-to-noise ratio (tSNR), framewise displacement (FD) and overall brain activity (z-score) during the entire scanning sessions using Matlab based scripts (Fig. 3C). In general, tSNR of the brain was higher in cortical regions (tSNR: 25 - 35) and gradually decreasing toward the subcortical area (tSNR: 15 - 25) due to the distance from the coil (Fig. 3C-i). To assess the motion induced artifacts, we set the threshold at 0.1 mm, and we did not observe any apparent motion across the fMRI time-series (Fig. 3C-ii). Furthermore, we did not observe any aberrant brain activation due to stimulation in awake state (Fig. 3C-ii). Altogether, these findings suggest the efficacy of our 3D printed cradle that can potentially be used for head-fixed awake fMRI experiments in rodents.

### fMRI data processing

#### Pre-processing

fMRI data was preprocessed using Statistical Parametric Mapping (SPM12, http://www.fil.ion.ucl.ac.uk/spm/) for MATLAB (MathWorks, USA) as described in a former report^18^ and DiffeoMap (www.mristudio.org).

#### Step 1-motion correction

Motion related artifacts from the raw data (Suppl. Fig. 4A) were corrected by realigning 300 fMRI volumes to the first scan using a least squares approach with a 6 parameters rigid body spatial transformation (Suppl. Fig. 4B).

#### Step 2-normalization

For group level analysis, fMRI data was co-registered to the respective T2 image volume using normalized mutual information approach, a 4th degree B-Spline interpolation and a 6-parameter rigid body transformation (3 parameters for translation and 3 parameters for rotation). fMRI and the anatomical data were then transferred into a template space using automated image registration (AIR) followed by large deformation diffeomorphic metric mapping (LDDMM) transformation implemented in DiffeoMap (^28^) (Suppl. Fig. 4C).

#### Step 3-smoothing

We applied a Gaussian smoothing with a kernel of full width at half maximum (FWHM) of 0.4 × 0.4 × 1 mm^3^ to all normalized fMRI datasets (Suppl. Fig. 2D). Step 4-segmentation: We further mapped the fMRI datasets into an atlas space (^16^) following the image registration procedures described in step 2 and segmented mouse brain into different structural labels (Suppl. Fig. 4E).

rs-fMRI datasets were temporally band-pass filtered (0.01 – 0.1 Hz) to capture only the low frequency fluctuations (LFFs) and further linearly detrended. Global signal regression was not applied to avoid false-positive induction of anti-correlation in the brain (^29^). A graphical illustration of the data preprocessing steps has been provided in the supplementary information (Sup. Fig. 4)

#### Post-processing

To justify the efficacy of the cradle in rs-fMRI experiments, we investigated whether the brain exhibits resting-state functional fingerprints during awake state using seed correlation analysis (SCA). Using the atlas (^16^), we extracted bi-lateral retrosplenial area (RSP) – the hub region of the default mode network (DMN) (^30^) and used as seed region for functional connectivity mapping in awake mouse brain. We computed the correlation coefficients (two-tailed *t*-test, *p* < 0.001) between the seed region and the averaged time series of the remaining whole brain and converted to *z* values using Fisher’s *r*-to-*z* transformation (^17,18,27^).

Functional clusters due to non-tactile stimulation were mapped using group independent component analysis (ICA) (^17,18,27^). Using the Matlab based toolbox GIFT (group ICA of fMRI toolbox-v1.3i, www.nitrc.org/projects/gift) and Infomax algorithm, we decomposed the entire BOLD data set into spatially independent components (ICs) without any hypothesis paradigms (^31^). Resulting mean functional clusters were displayed as spatial color-coded z-scores onto template T2 image (threshold |*z*| > 3, *p* < 0.00135) where the color coding represents the dependence of the time course in each voxel compared with the mean time course of the respective component in arbitrary units.

### Multimodal imaging in anesthetized and awake mice

We further demonstrated the adaptability of our animal conditioning set-up to other preclinical imaging systems, such as, computed tomography (CT), positron emission tomography (PET), and PET/CT. The animal was initially anesthetized using isoflurane in a specific chamber and stereotaxically fixated in the cradle using tooth bar, ear bars, and nose cone. The nose cone was attached to an isoflurane delivery system using a single 2 mm diameter plastic tube.

Animal was breathing through the experiment under a steady stream of isoflurane (0.5-2%) mixed with air.

Both anesthetized and awake animals went through MRI imaging as follows: FLASH sequence to verify the centering, then T2-weighted RARE sequence with parameters as follows: TR=2516 ms, TE=35 ms, Repetitions = 20, Turbo-factor=8, FOV=30 mm x 9 mm, Matrix size=256 × 256 resulting in 117 µm x 35 µm in-plane resolution with a total of 19 slices of 800 µm thickness. Scans were averaged over 2 times to maximize SNR resulting in a total imaging time of 26 min and 50 seconds.

The plastic tube attached to the nose-cone was then detached from the isoflurane delivery system and the cradle, holding the mouse still under anesthesia, was moved quickly to the next room where the Inveon (Siemens, Munich, Germany) imaging system was located. The nose-cone tube is then attached to the isoflurane delivery system in this room as quickly as possible making sure that the animal does not wake up. The 3D printed platform is attached to the Inveon animal bed using the specific 3D adaptor, which is attached to the cradle holding the animal on one side and the Inveon system on the other.

The animal is then inserted into the scanner and went through PET/CT dynamic imaging. Both animals were injected with Fludeoxyglucose (FDG) dose (200 mCi in 200ul volume) dynamically using an automatic pump at a rate of 120ul/min for the PET scan. In the case of anesthetized mouse, the body temperature of the animals was maintained by using an air heat-supply during the scan. Both anesthetized and awake mice were imaged using the same PET/CT acquisition parameters. PET acquisition parameters were: Emission= 1800 sec, axial length = 127 mm. CT acquisition parameters were: attenuation scan= 528 sec, effective pixel size= 99.89 µm, transaxial FOV=51.15 mm, axial FOV=79.92 mm, exposure settings (80 Kv, 500 µA, filter 0.5 mm, exposure 180 ms).

### Analysis of PET/CT scan data

Inveon Research Workplace (IRW) software package (Siemens) was used to analyze the PET/CT data. We first co-registered the CT and PET data using anatomical landmarks. Then brain region of interest (ROI) were defined using the geometric tools. Next, kinetic and quantitative analysis were performed on the brain ROI by measuring of the Standard Uptake Value of the FDG.

## SUPPLEMENTARY FIGURES

**Supplementary Figure 1.**
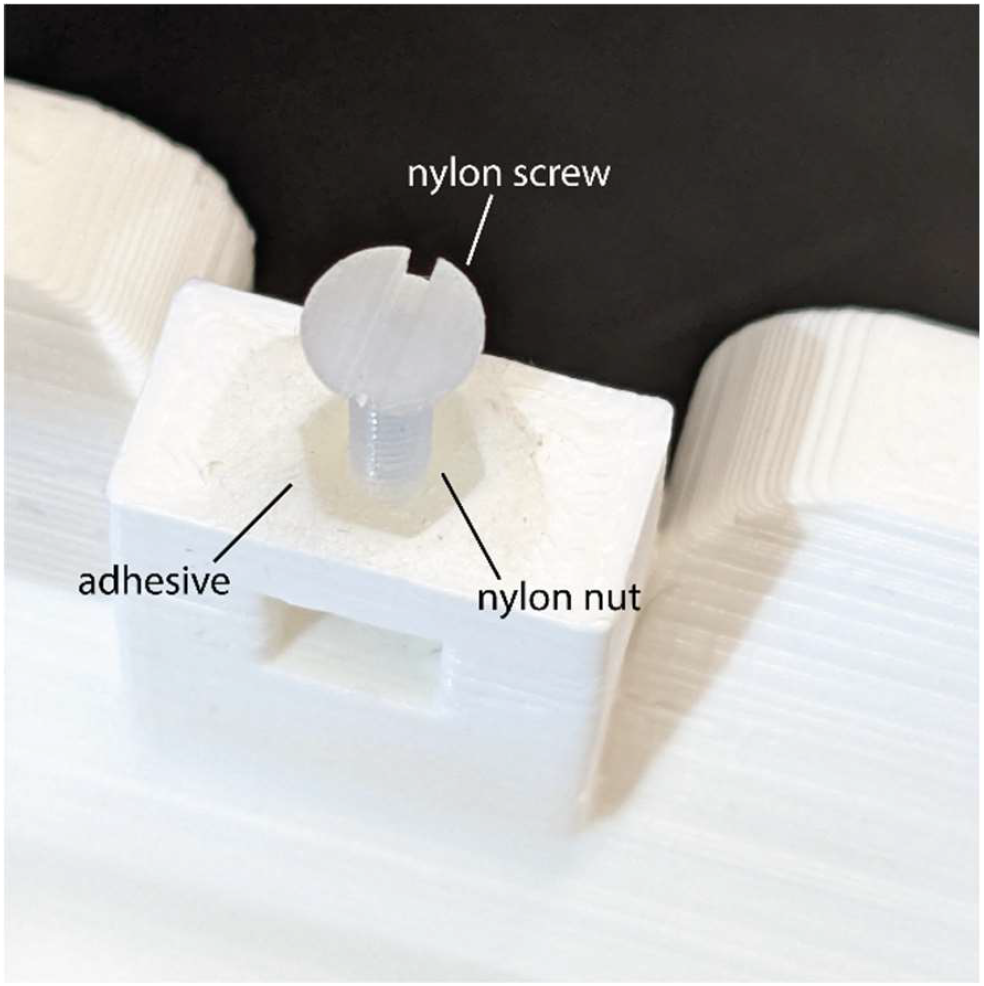
Tap-free screw mechanism. Instead of tapping nylon nuts are glued to the printed parts in designated places (to host the adapting screws) using adhesive materials.

**Supplementary Figure 2.**
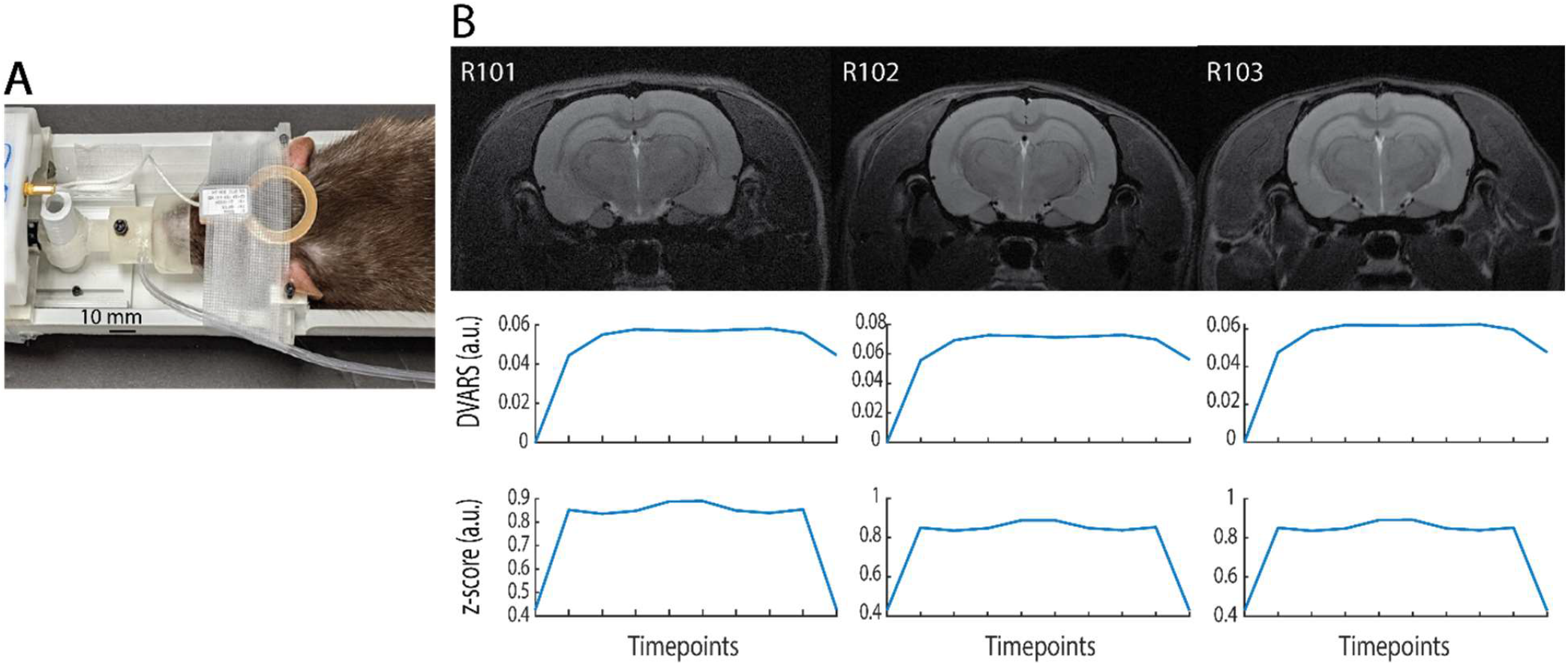
Motion assessment during the MR imaging. The consistency of high values on Z-score and DVARS measures over 10 repetitive images across animals (related to same data illustrated in Fig. 2B) confirm resilience of the animal preparation using the set-up to motion artifacts.

**Supplementary Figure 3.**
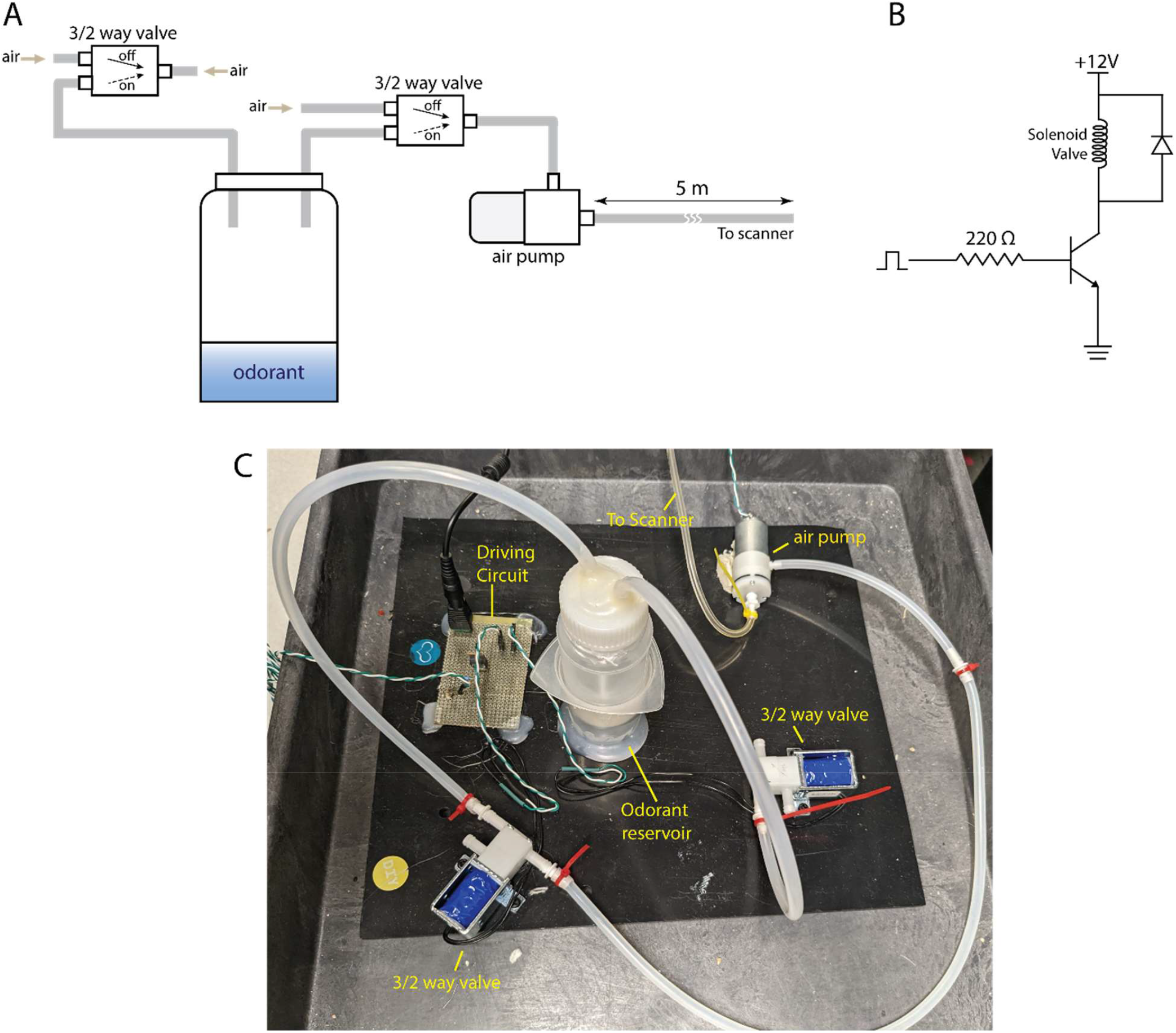
Odor delivery system. **A)** Graphical representation of the overall system design. **B)** Schematic of the electrical circuit for driving the valves. The air pump is powered by a DC supply to be kept continuously ON (circuit not shown). **C)** Picture of odor delivery system used in our experiments.

**Supplementary Figure 4.**
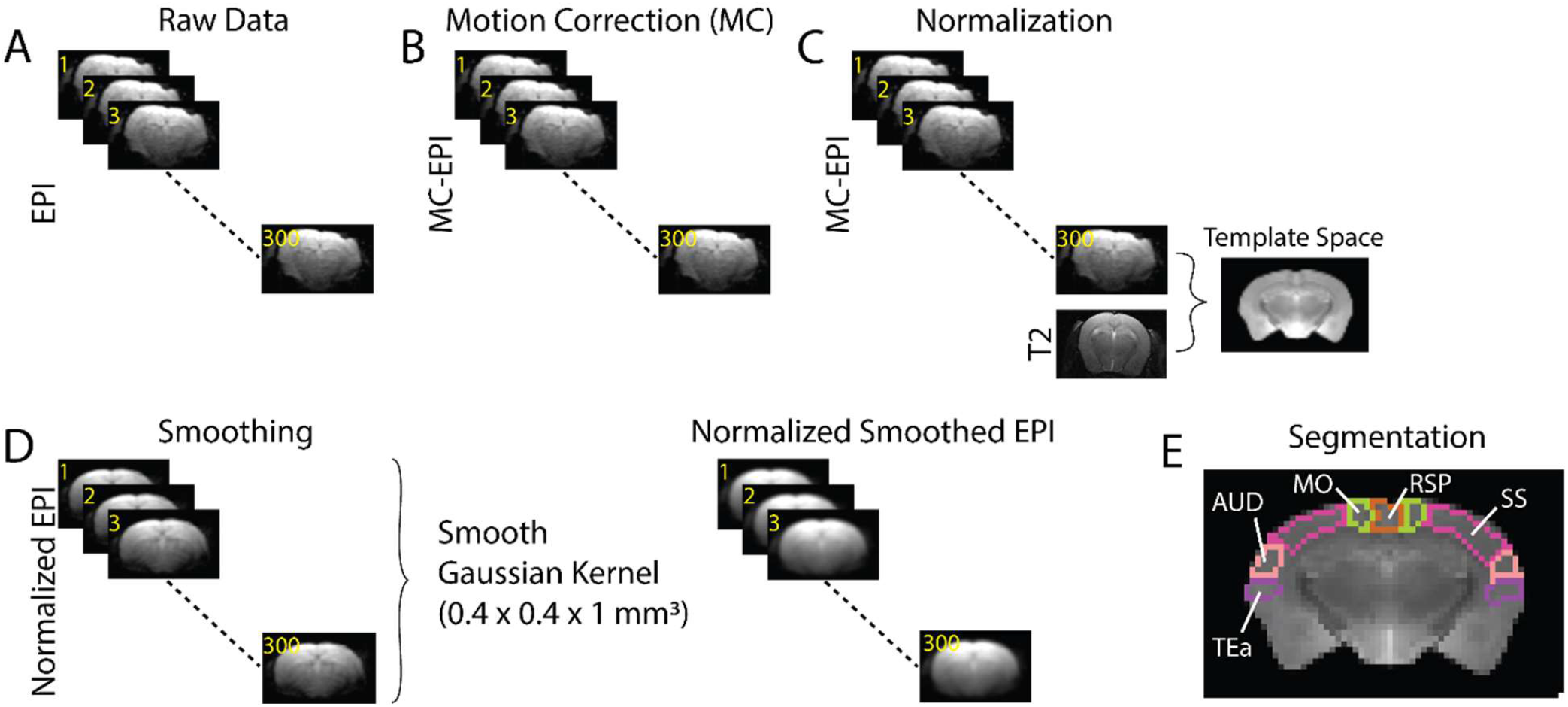
Process flow for analysis of the fMRI data. **A)** Rawdata containing 300 EPI datasets from a specific scan. **B)** Motion correction; realignment of the 300 EPI volumes to the first one. **C)** Normalization; Motion corrected EPI and the corresponding anatomical T2 images were coregistered into a template space using rigid AIR and Large Deformation Diffeomorphic Metric Mapping (LDDMM) transformation. **D)** Normalized EPI data were smoothed with a Gaussian kernel of full width at half maximum of 0.4 × 0.4 × 1 mm^3^. **E)** Segmentation; whole brain was parcellated using an in-house built mouse brain atlas.

## Notes

### Competing Interest Statement

The authors have declared no competing interest.

